# Task Demands Predict a Dynamic Switch in the Content of Awake Hippocampal Replay

**DOI:** 10.1101/172098

**Authors:** H. Freyja Ólafsdóttir, Francis Carpenter, Caswell Barry

**Affiliations:** Research Department of Cell and Developmental Biology, UCL, Gower Street, London. WC1E 6BT. UK.; Institute of Neurology, UCL, Queen Square, London. WC1N 3BQ. UK.

**Keywords:** place cell, grid cell, entorhinal cortex, hippocampus, replay, planning, memory consolidation, navigation

## Abstract

Reactivation of hippocampal place cell sequences during behavioural immobility and rest has been linked with both memory consolidation and navigational planning. Yet it remains to be investigated whether these functions are temporally segregated; occurring during different behavioural states. During a self-paced spatial task, awake hippocampal replay occurring immediately before movement towards a reward location, or just after arrival at a reward location, preferentially involved cells consistent with the current trajectory. In contrast, during periods of extended immobility, no such biases were evident. Notably, the occurrence of task-focused reactivations predicted the accuracy of subsequent spatial decisions. Additionally, during immobility but not periods preceding or succeeding movement, grid cells in deep layers of entorhinal cortex replayed coherently with the hippocampus. Thus, hippocampal reactivations dynamically and abruptly switch operational mode in response to task demands – plausibly moving from a state favouring navigational planning to one geared towards memory consolidation.

## Introduction

Prominent theories of hippocampal function place it at the centre of networks supporting memory and navigation(O’Keefe and Nadel, 1978; Scoville and Milner, 1957). The principal cell of the hippocampus is the ‘place cell’, whose activity during locomotion encodes the animal’s self-location via spatially localised firing fields (‘place fields’)(O’Keefe and Dostrovsky, 1971). However, during non-REM sleep and pauses in locomotion, when sharp-wave ripple complexes (SWRs) transiently dominate the hippocampal local field potential (LFP)(Buzsaki et al., 1992; O’Keefe and Nadel, 1978), place cell activity de-couples from the animal’s current location, re-activating past or future spatial trajectories (‘replay’)(Foster and Wilson, 2006; Lee and Wilson, 2002; Wilson and McNaughton, 1994).

At the time of discovery, replay was proposed as the mechanism supporting systems-level memory consolidation (Wilson and McNaughton, 1994); the process by which memories become less susceptible to hippocampal damage (Marr, 1971; Scoville and Milner, 1957). Consistent with this hypothesis, replay typically reflects recent experiences, particularly novel ones (Cheng and Frank, 2008; Foster and Wilson, 2006; O’Neill et al., 2008; van de Ven et al., 2016), is dependent on the NMDA receptor (Dupret et al., 2010; Silva et al., 2015), and is associated with cortical reactivations (Ji and Wilson, 2007; Rothschild et al., 2016; Wierzynski et al., 2009). Indeed, cortical replay has been found to temporally lag the hippocampus (Olafsdottir et al., 2016a; Rothschild et al., 2017), suggestive of information flow from the hippocampus to the cortex (Olafsdottir et al., 2016b; Rothschild et al., 2017). Furthermore, various studies have shown that cortical LFP patterns associated with sleep, such as delta waves (Maingret et al., 2016; Mednick et al., 2013) and spindles(Johnson et al., 2010), are temporally coordinated with SWRs (Battaglia et al., 2004; Peyrache et al., 2011; Sirota et al., 2003), and have indicated that cortico-hippocampal dialogue may be important for learning (Maingret et al., 2016). More generally, SWRs originate in the hippocampus(Buzsaki, 2015; Suzuki and Smith, 1985), propagate into the cortex(Chrobak and Buzsaki, 1994, 1996), and occur at a greater rate after learning (Eschenko et al., 2008). Elimination of SWRs during rest impairs spatial learning (Ego-Stengel and Wilson, 2010; Girardeau et al., 2009a) and conversely hippocampal reactivation during rest enhances learning (de Lavilleon et al., 2015; Rasch et al., 2007).

Nevertheless, it is now apparent replay and the roles attributed to it are more diverse than first thought. While the role of replay during non-REM sleep (‘offline’) in consolidation is well supported, the purpose of awake replay (‘online’) is less clear. On one hand, online replay is modulated by environmental novelty (Cheng and Frank, 2008; Foster and Wilson, 2006) as well as changes in reward (Ambrose et al., 2016; Singer and Frank, 2009), and interference with online SWRs impairs the acquisition of spatial tasks (Jadhav et al., 2012); suggestive of a role in learning, if not also consolidation. However, online replay has also been linked with spatial planning and navigation (Foster and Knierim, 2012a; Pfeiffer and Foster, 2013; Samsonovich and Ascoli, 2005). Theoretical propositions suggest replay as a mechanism for exploring potential routes or extracting goal-directed heading vectors (Bush et al., 2015; Erdem and Hasselmo, 2012; Erdem and Hasselmo, 2014; Gupta et al., 2010). A view supported experimentally by results demonstrating preferential replay of goal-oriented trajectories (Pfeiffer and Foster, 2013; Singer et al., 2013).

This complexity with regards to the function of online replay is mirrored by distinctions in its form. For example, the sequence in which place fields are reactivated during replay can recapitulate their order on the track (‘forward’ replay)(Diba and Buzsaki, 2007a, b; Lee and Wilson, 2002; Skaggs and McNaughton, 1996) or invert it (‘reverse’ replay)(Foster and Wilson, 2006); a dichotomy that also exists for offline replay (Olafsdottir et al., 2016b; Wikenheiser and Redish, 2013). While forward replay was initially linked with movement initiation and reverse with goal-arrival (Diba and Buzsaki, 2007a), that distinction is now less secure (e.g.(Gupta et al., 2010)), and it appears reverse replay may play a role in reinforcement learning (Ambrose et al., 2016; Foster and Knierim, 2012b). Furthermore, although online replay typically encompasses locations close to the animal’s current location (Davidson et al., 2009; Dupret et al., 2010; O’Neill et al., 2006; Pfeiffer and Foster, 2013) it can also be remote, representing distant portions of the current environment not related to current goals (Davidson et al., 2009; Gupta et al., 2010), a portion of an environment being avoided(Wu et al., 2017), or even entirely distinct enclosures (Jackson et al., 2006; Karlsson and Frank, 2009).

Thus, although navigation is sometimes associated with forward replay of proximal locations, perhaps supporting goal-directed navigation, equally often online replay is not navigationally relevant, depicting remote location, away from important goals. The factors that govern the switch between these two forms of replay remains unclear; what influences the content of reactivations? Could task demands dictate whether replay is conducted for the purpose of planning or consolidation? If so, does the occurrence of navigationally relevant replay contributes to accurate spatial decisions?

Here, we analyse place cell replay occurring during periods of immobility interspersing a self-paced spatial decision task. We find that after pausing, rats transition rapidly from a state in which they preferentially exhibit replay associated with the ongoing task, to a ‘disengaged’ state in which replay was directed towards remote sections of the track and also incorporated grid cells from deep layers of the entorhinal cortex. Subsequently, reinstatement of movement was prefixed by a switch back to an ‘engaged’ state; characterised by forward replay of proximate sections of the track without coherent grid activity. Finally, we observed that reactivations occurring before animals made errors on the task did not exhibit engaged-like replay and errors could be predicted on the basis of the replay content. These results suggest that online replay supports both consolidation and spatial planning, providing the first evidence that switching between these states occurs dynamically according to task demands and that when animals are engaged in a task the replay content can be important for accurate spatial behaviour.

## Results

We recorded CA1 place cells (21-72 cells/session, Figure 1C, Figure S1, Table 1) over 1-5 days in eight rats whilst they performed a spatial decision task (see Materials and Methods, Figure 1A). Animals were required to complete laps on an elevated 6m Z-track. Specifically, rats began each session at the end of arm1 and ran to the first corner (between arm1 and arm2) where they stopped in order to receive a food reward. Following this, they ran to the second corner (between arm2 and arm3) and then on to the end of arm 3, being rewarded at the corner and arm end. They then ran back to the start of arm1 in a similar fashion (mean number of laps = 22 (SD = 4.82), mean pause duration= 10.71s (SD = 16.49s, Figure S2)). Note, only correct turns at the corners would result in food reward at the next stop. Runs going from arm1 to arm3 were labelled ‘outbound’ and runs going back from arm3 to arm1 ‘inbound’. With experience, animals became increasingly fluent at the task, making fewer wrong turns at the corners (error vs. day, r = -0.61, p = 0.0016 Figure 1B) and taking less time to complete laps (lap duration vs day, r = -0.46, p = 0.0066).

**Figure 1.**
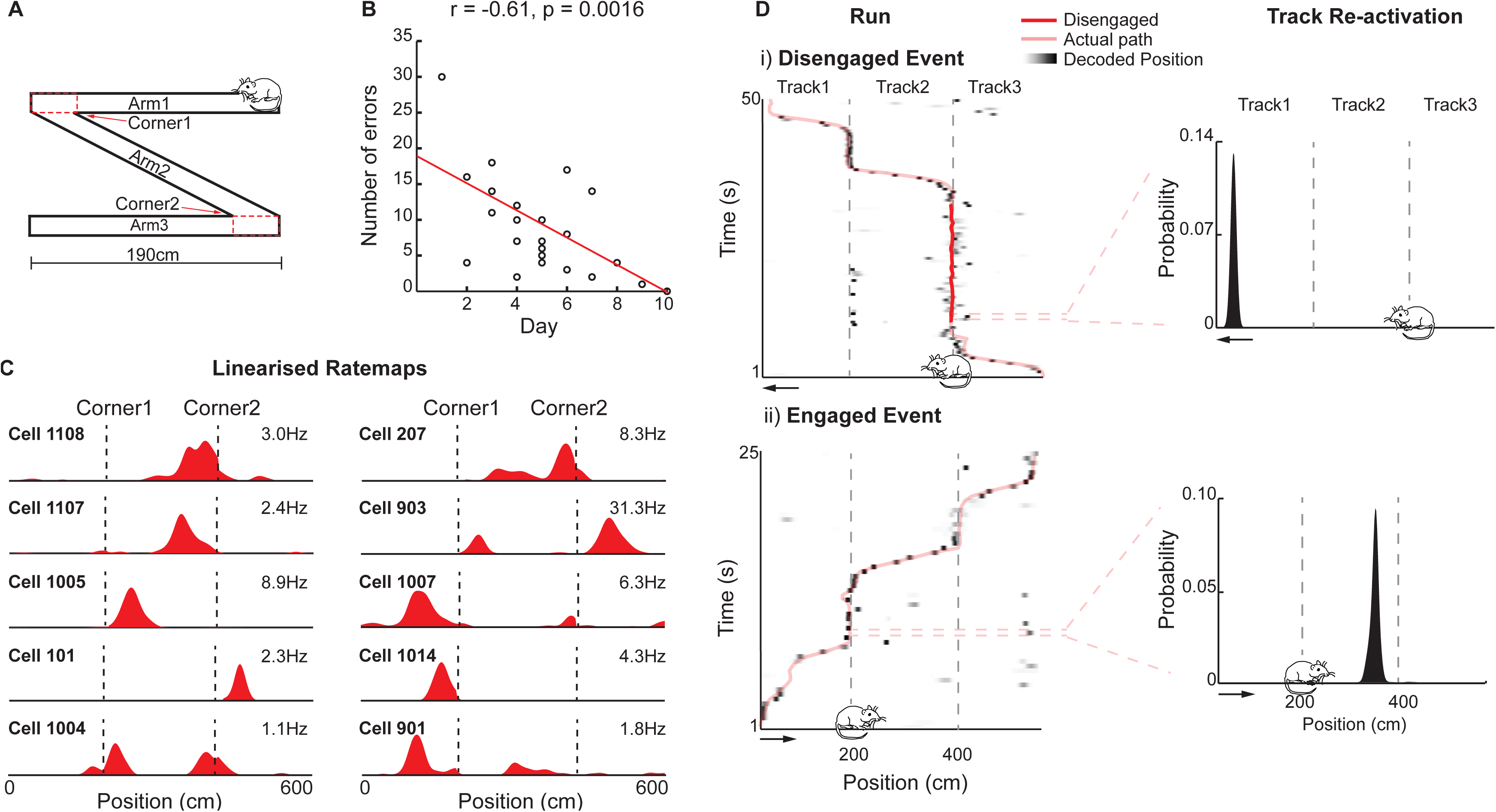
Hippocampal Reactivations during Task Engagement and Disengagement. (**A**) Rats completed laps on the Z-track. (**B**) Number of errors per session vs. experimental day, title shows Pearson’s r. (**C**) Representative linearised outbound place cell ratemaps for 10 CA1 place cells, x-axis: track position (cm), y-axis: firing rate scaled to each cell’s peak rate (shown above the ratemap). (**D**) Representative disengaged and engaged reactivation events. i) Left: an example path (pink) superimposed on the posterior probability matrix over position, darker shades indicating higher probability. Position of cartoon rat indicates the animal’s location and heading direction during the event. Disengaged periods highlighted in red, x-axis: track position (cm), y-axis: time (s) (bin size: 500ms). Right: Position decoding of a reactivation event, x-axis: track position (cm), y-axis: probability of position. ii) Same as i) but for an engaged reactivation event. See also Figures S1-S3.

Corner stops were divided into two temporal sections: ‘engaged’, including periods when the animal had just arrived at the corner (< 5s) and just before movement onto then next track was reinstated (< 5s, Figure 1D(ii)), and ‘disengaged’, the time interleaving these two periods (Figure 1D (i)). We analysed reactivation events occurring while animals were at the corners. Reactivations were identified based on increases in multi-unit place cell activity (see method for details) and were limited to periods when the animals’ speed remained below 3cm/s. Place cell activity during reactivations was analysed using a Bayesian decoding approach(Davidson et al., 2009; Olafsdottir et al., 2016b; Zhang et al., 1998) to calculate the probability of an animal’s location on the track given the observed activity (Figure 1D, Figure S3). In the first instance, to understand which sections of the track were represented, we simply summed the probability distribution over each arm, identifying the one which was most strongly reactivated (Figure 1D (i-ii)). Note, because place cell activity is highly directional on linear tracks (McNaughton et al., 1983), inbound and outbound runs on each arm were treated separately (mean Pearson correlation between inbound and outbound ratemaps = -0.0032 (SD = 0.28)). Thus, it was possible to identify reactivation of each of the three arms, in either the inbound or outbound direction.

We identified a total of 1415 engaged and 3010 disengaged reactivation events. No differences were found between engaged events occurring at the start and end of pauses (Figure S4); as such, these periods were combined for all subsequent analyses. Engaged events showed a robust bias to reactivate spatial firing ‘congruent’ with an animal’s current direction of travel (62.26% vs chance p < 0.0001, chance derived from shuffling cell IDs, figure 2A(i-ii)&B). Hence, during inbound runs, inbound representations were more likely to be reactivated and vice versa. Disengaged events showed no such bias, being equally likely to activate congruent or ‘incongruent’ representations (52.19% vs chance, p = 0.39, engaged vs disengaged, p < 0.0001, Figure 2B). Moreover, engaged events also showed a strong preference to reactivate either of the tracks adjacent to the animal’s current position, the ‘local’ arms, as oppose to the ‘remote’ arm (80.70% local reactivation vs chance, p < 0.0001, figure2A(iii-iv) & C). Again, this bias was not present in disengaged events, where the proportion of events where the local arms were reactivated relative to the remote arm was not different to chance (71.42% vs chance, p = 0.31; engaged vs disengaged, p < 0.0001). Importantly, the results could not be explained by differences in ripple power (Figure S5) or movement speed (Figure S6) during the two types of events. Although disengaged events were characterised by lower theta-band power than engaged events (log(theta power/delta power) during engaged = 0.31 (SD = 0.41), disengaged = 0.23 (SD = 0.39), p < 0.0001, Figure S5) results were unchanged by the exclusion of events with high theta power (i.e. analysis limited to events with theta power < 1SD below mean power during movement, Figure S7). Thus, reactivation events occurring while animals were engaged in the task were more likely to incorporate place cells immediately relevant to that task; representing adjacent sections of the track and congruent with the current direction of travel.

**Figure 2.**
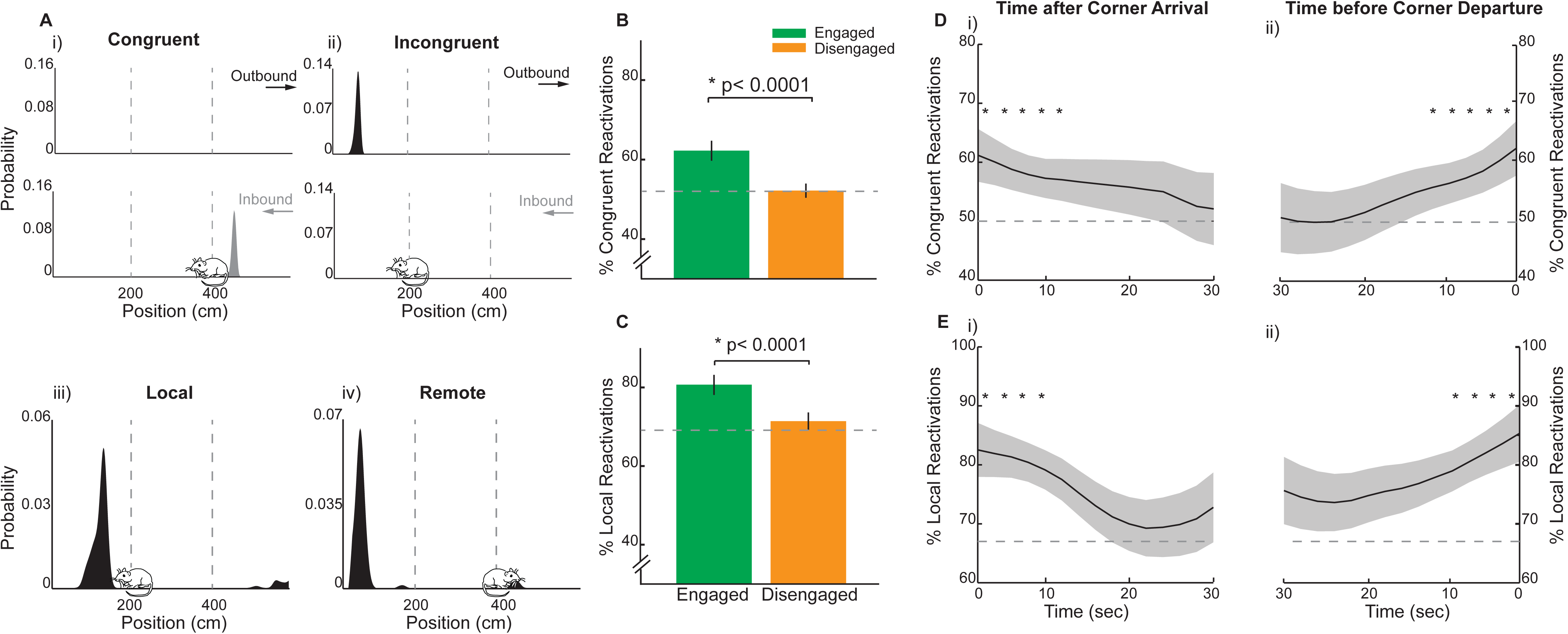
Track Reactivations are Modulated by Task Engagement. (**A**) Representative position decoding for a congruent (i), incongruent (ii), local (iii) and remote (iv) reactivation event. Upper panels: Decoding was done for outbound (top) and inbound (bottom) runs separately. Reactivation events decoding to a run consistent with the animal’s current trajectory were categorised as ‘congruent’ (left). Lower panels: events reactivating positions on the arms immediately ahead or behind the animal were categorised as ‘local’ (iii), while events reactivating positions on the remote arm were categorised as ‘remote’ (iv). Position of animal during event indicated by location of cartoon rat, x-axis: track position (cm), y-axis: probability of position. (**B**) Proportion of engaged (green) and disengaged (amber) reactivation events categorised as congruent. Error bars indicated 95% confidence interval based on bootstrapped data. (**C**) Same as (B) but for local reactivations. (**D**) Proportion of congruent reactivations as a function of time after corner arrival (i) and before corner departure (ii). Shaded areas show 1 standard deviation of bootstrapped data, x-axis: time (s) (bin size = 2sec), y-axis: proportion congruent reactivations. (**E**) Same as (D) but for local reactivations. ^*^p< 0.05 vs chance. See also Figures S4-S7.

To better understand the time course governing the transition between the engaged and disengaged states, we examined how the proportion of local and congruent reactivations varied as a function of the time since arrival at, as well as departure from, the corners. Specifically, while animals were stationary (<3cm/s) at the corners, we calculated the proportion of congruent/local events in 5s windows (advancing in 2.5s increments) and assessed at what time point the proportions were no longer significantly above chance. Following corner arrival, the proportion of congruent events remained above chance for the first 12.5s (Figure 2D(i)). A bias for congruent re-activations was also present for the last 12.5s preceding corner departure (Figure 2D(ii)). The analysis of local reactivations revealed a similar pattern; namely, the proportion of events classified as local remained above chance for the first 10s after corner arrival (Figure 2E(i)) and for the last 10s prior to exit (Figure 2E(ii)). Thus, for the first and last 10-15s of stopping periods hippocampal reactivations remained in an ‘engaged’ state, after which they sharply transitioned to a ‘disengaged’ state.

Subsequently, we examined the trajectories encoded during reactivation events. A Bayesian approach was applied to calculate, for each 10ms bin within an event, the probability distribution over track position (Zhang et al., 1998). We then applied a line fitting procedure(Davidson et al., 2009; Olafsdottir et al., 2016b) to the resulting posterior probability matrix to identify possible replay trajectories (Figure 3A(i-ii)). Again, events were classified as either outbound or inbound depending on which produced the higher best line-fit. Putative events whose rank against their own spatial shuffle distribution exceeded the 97.5^th^ percentile, and which occurred while the animal was stationary (<3cm/s), were classified as replay events (Figure S8 Table S1).

**Figure 3.**
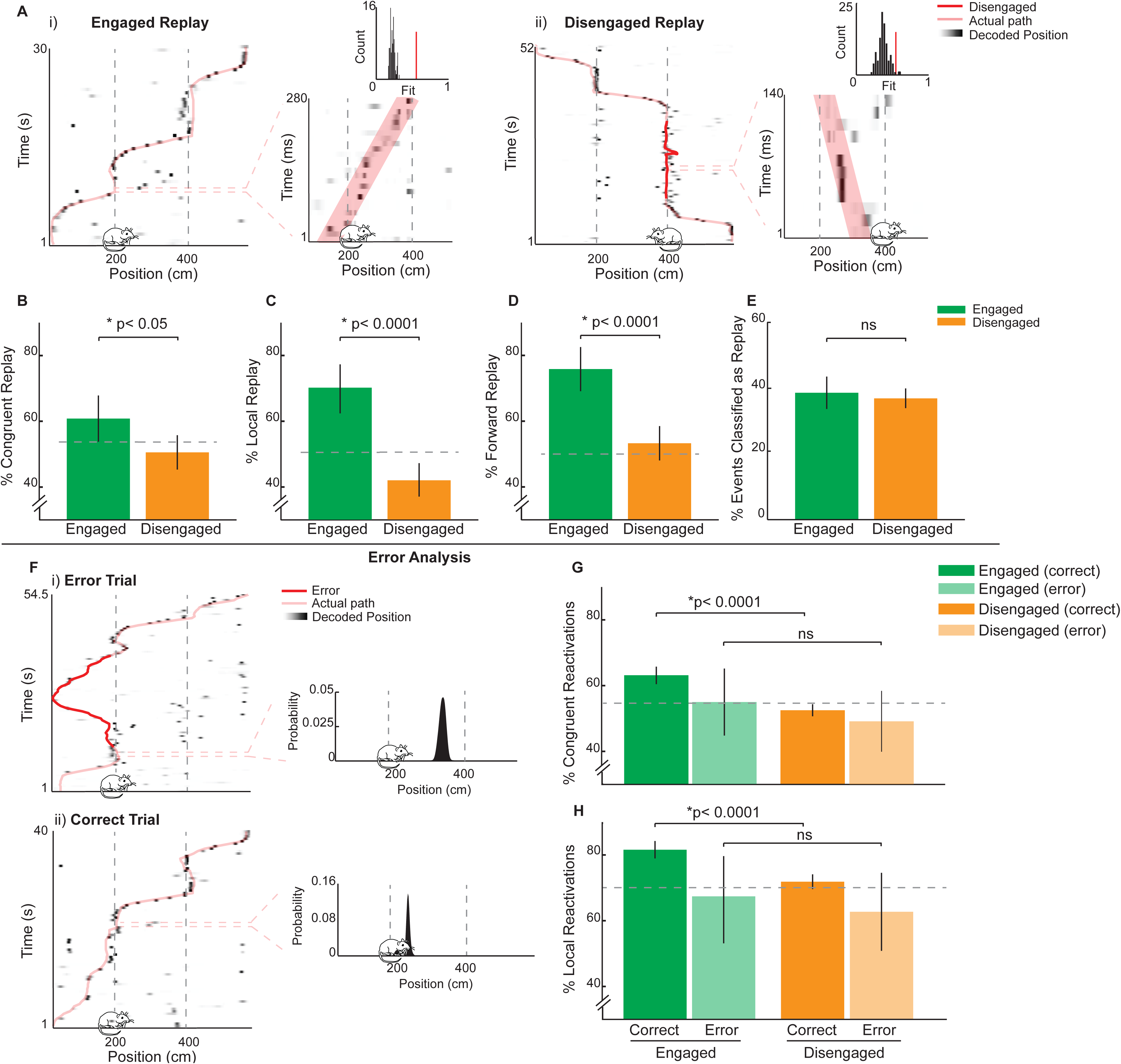
Reactivations During Engaged Periods Prefigure Subsequent Behavioural Performance. (**A**)Representative engaged(i) and disengaged(ii) replay events. i) Left: path (pink) superimposed on position decoding, darker shades indicating higher probability of position. Cartoon rat indicates animal’s position during event, x-axis: track position (cm), y-axis: time (s) (bin size = 500ms). Right: Position decoding during a replay event, best fit trajectory superimposed in pink. x-axis: track position (cm), y-axis: time (ms) (bin size = 10ms). Inset: replay event best fit trajectory shuffle distribution, red bar indicates best fit trajectory score of original event. ii) same as i) but for a disengaged event. Disengaged period highlighted in red. (**B**) Proportion of engaged (green) and disengaged (amber) replay events categorised as congruent. Error bars show 95% confidence interval based on bootstrapped data. (**C-D**) Same as (B) but for local and forward replay. (**E**) Proportion of putative engaged (green) and disengaged (amber) events categorised as replay events. Error bars same as in (B-D). (**F**) Representative reactivation events preceding error and correct spatial choices. i) Left: rat’s path (pink) superimposed on position decoding, darker shades indicating higher probability of position. Erroneous portion of the path is shown in red. x-axis position on the track, y-axis time (s), bin size = 500ms. Right: position decoding of reactivation event. Rat location indicated by position of cartoon rat, x-axis: track position (cm), y-axis: probability of position. ii) Same i) but for a reactivation event preceding a correct spatial choice. (**G**) Proportion of engaged (green) and disengaged (amber) events classified as congruent. Darker bars show proportions for events preceding a correct spatial choice, lighter bars indicate errors. Error bars indicate 95% confidence interval based on bootstrapped data. Interaction between error/correct and engaged/disengaged was not significant (p=0.19, difference engaged correct vs. engaged error = 9.7%, difference disengaged error vs. disengaged correct = 3.4%) (**H**) Same as (G) but for local reactivations. Again, interaction was not significant (p=0.14, difference engaged correct vs. engaged error = 15.0%, difference disengaged error vs. disengaged correct = 7.6%). See also figures S8-S14.

We identified a total of 149 engaged replay events and 364 disengaged replay events. As before, we saw that engaged replay events preferentially reactivated spatial representations congruent with an animal’s current direction of travel (60.82% vs chance, chance derived from shuffling of engaged and disengaged events, p = 0.024; engaged vs disengaged, p = 0.013, Figure 3B). Disengaged events exhibited no bias, being equally likely to represent trajectories moving in either direction (50.50% vs chance, p = 0.89). Furthermore, a clear majority of engaged events replayed trajectories close to the animal (median replay position <60cm from animal, 70.21% vs chance, p < 0.001, mean distance from animal at start of event = 69.32cm (SD = 80.78cm)) whereas disengaged events showed a preference for replaying trajectories remote to the animal (42.03% vs chance, p = 0.0006; engaged vs disengaged, p < 0.0001, mean distance from animal at start of event = 134.48cm (SD = 119.17cm) p < 0.0001, Figure 3C, Figure S9). Moreover, we classified replay events as ‘forward’ or ‘reverse’ based on the slope of the best fit line and found the majority of engaged replay events were forward events (75.8% vs chance (50%), p < 0.0001) while disengaged periods did not show a preference for forward or reverse replay (53.3% vs chance, p = 0.093, engaged vs disengaged, p < 0.0001, Figure 3D). Importantly, these differences could not be explained by the robustness of the replayed trajectories; although we identified a higher number of disengaged replay events than engaged events, the percentage of putative events qualifying as replay events for the two categories was not different (36.88% vs 38.60%, p = 0.28 Figure 3E). It is unlikely that the observed difference between engaged and disengaged periods result from theta sequences (Foster and Wilson, 2006; Gupta et al., 2012; Johnson and Redish, 2007) or phase precession(O’Keefe and Recce, 1993): theta power during engaged and disengaged replay events did not differ (log(theta/delta) power during engaged = 0.11, disengaged = 0.13, p = 0.65, Figure S10) and results were unchanged when replay trajectories were required to exceed 2m/s speed (Figure S11) or when they were limited to events that co-occurred with SWRs (Figure S12). To sum, during periods of task engagement, replay preferentially depicted task relevant place cell sequences: trajectories congruent with an animal’s current direction of travel, that were close to its current position, and which maintained the order in which cells were activated during experience. Conversely, during disengaged periods, the replay trajectories were less biased; they were equally likely to depict motion in either direction regardless of the animal’s heading, showed no preference for forward and reverse sequences, and were more focused on distant regions of the track.

Medial entorhinal (MEC) grid cells are also known to exhibit replay (O’Neill et al., 2017; Olafsdottir et al., 2016b). During rest, grid cells from the deep layers of the MEC – the primary cortical projection target of the hippocampus - replay coherently with place cells but do so with a slight delay (11ms)(Olafsdottir et al., 2016b). Conversely, grid cells from the superficial layers of the MEC – afferent to the hippocampus – replay trajectories independently of the hippocampus, an effect that is more pronounced during task engagement than rest(O’Neill et al., 2017). Plausibly, the transmission of replayed trajectories from CA1 to the MEC during quiescence might support consolidation, whereas hippocampal-independent replay in superficial MEC is more likely to be linked with navigational planning(Bush et al., 2015) and need not engage deep MEC. To investigate this distinction, we analysed recordings made concurrently from deep MEC (V/VI) and CA1 in seven of the eight rats included in the study. During both engaged and disengaged periods we observed LFP coherence between the hippocampus and MEC in the theta (6-12Hz) and ripple (150-250Hz) bands (Figure S13). Specifically though, we found grid cells were coordinated with place cell replay during disengaged periods (grid-place coordination = 0.097 (SD = 0.08) vs chance p = 0.011) but not engaged periods (grid-place coordination = 0.074 (SD = 0.037) vs chance p = 0.75) the amount of coherence between the two periods being markedly different (engaged vs disengaged, p = 0.0035, Figure S13). As such, these results suggest a further functional distinction between replay occurring during engaged and disengaged periods; in the latter case, the engagement of cortical grid cells, would appear to provide a mechanism suitable to support systems-level consolidation.

To what extent is the content of engaged reactivations important for accurate task performance? To address this question, we analysed events preceding correct and incorrect turns separately (Figure 3F (i-ii)). Due to the relatively small number of errors (mean wrong turns per session = 8.88 (SD = 6.88)) analyses were limited to simple reactivation events. For events preceding correct turns we saw, as before, a clear distinction between engaged and disengaged reactivations (Figure 3G&H). Thus, engaged reactivations had a tendency to be congruent with, and local to, an animal’s position (62.79% congruent vs chance, p< 0.0001; 81.5% local vs chance, p < 0.0001), while disengaged reactivations exhibited no such bias (52.34% congruent vs chance, p=0.19; 71.76% local vs chance, p=0.12, engaged vs disengaged both ps < 0.0001). However, for events preceding incorrect turns, this distinction was absent. As expected, disengaged events exhibited no bias towards congruent or local reactivations (48.76% congruent vs chance, p=0.94; 62.71% local vs chance, p=0.89) but neither did, in this instance, engaged events (54.44% engaged, p=0.76 vs chance; 67.35% local vs chance, p=0.67; engaged vs disengaged both ps > 0.20). Indeed, the proportion of engaged local events preceding incorrect turns was significantly lower than that preceding correct turns (p < 0.0001), a marginally significant effect in the same direction was observed for congruent events (p = 0.062). However, due to the small number of error events (engaged error events = 90, disengaged error events = 121), we did not find a significant interaction between levels of task engagement and the accuracy of future decisions (based on bootstrapping difference scores for events preceding correct and erroneous turns, congruent
analysis p=0.19, local analysis p=0.14, Figure 3G&H). To understand if the observed difference was also manifest at the network level we examined how power in the ripple band of the LFP (150-250Hz) varied between events preceding correct and incorrect choices. Ripple power did not differ between disengaged events preceding correct and incorrect turns (correct = 1.73 × 10^−9^V^2^/Hz, error = 1.61 × 10^9^ V^2^/Hz, p=0.76, Figure S14). However, for engaged events alone, we saw that ripple power was reduced prior to errors (correct = 1.62 × 10^-9^ V^2^/Hz, error = 1.27 × 10^-9^ V^2^/Hz, p = 0.021, Figure S14). Again, we found the interaction analysis was in the right direction but did not reach statistical significance (p = 0.16). Thus, the content of engaged reactivations predicts the accuracy of future decisions; prior to making erroneous spatial decisions, reactivations were less task-focused and were accompanied by a reduction in ripple power.

Finally, to corroborate the suggestion that task-relevant reactivations are important for accurate spatial behaviour, we trained a binary classification decision tree (Kohavi and Quinlan, 2002; Rokach and Maimon, 2008) to predict the occurrence of errors based on all three of the measures we had obtained for the preceding reactivations (see Star Methods). Specifically, the decision tree was trained with data specifying for each event if it was local to the animal, congruent with the current direction of travel, as well as its mean power in the ripple band. Training and test data was partitioned using a 10-fold cross validation scheme; on each of 10 iterations the model was fit to 90% of the data and tested on the remaining 10%, using different portions of the data on each fold. Trees trained on reactivations occurring during engaged periods predicted the outcome of subsequent decisions correctly 62.8% of the time (p=0.001 compared to 1000 iterations of the model fit to shuffled data, mean shuffle prediction accuracy = 49.6%). Similarly, training using just the locality and congruence of events also allowed for better than chance prediction performance (60.3% accuracy, p=0.02 vs shuffle). Note, it is likely that the classification accuracy is not higher, precisely because reactivations occurring before errors are heterogeneous. In contrast to the engaged periods, disengaged reactivations were uninformative about the accuracy of subsequent decisions (55.2% decision accuracy, p=0.09 compared to 1000 iterations of the model fit to shuffled data, mean shuffle prediction accuracy = 49.8%). Thus, when considered in toto, the content of reactivations during periods of task engagement predicts whether the animal’s future choice will be accurate.

## Discussion

We have described findings showing a clear distinction between replay occurring while animals were engaged in a spatial task vs. periods in which they were simply resting on the track. When animals were preparing to make a spatial decision and initiate movement, or when they had just completed a trajectory, replay tended to be focused on its immediate environment; reactivating trajectories consistent with the animal’s position and heading, in a forward direction. Conversely, during prolonged stops at the corners of the track, replay trajectories were distributed across the apparatus, were equally likely to propagate in a forward or reverse direction, and incorporated entorhinal grid cells from the deep layers of MEC. The transition between these states occurred relatively rapidly, being task focused for 10-15s before and after movement, and the presence of task-focused reactivations during periods when animals were engaged with the task was found to predict the accuracy of their decisions. In sum, these results show the content of awake hippocampal reactivations can transition predictably and dynamically in accordance with task demands and that the content may determine the accuracy of spatial behaviour.

Replay, particularly online replay, has been mooted to support a range of neurobiological phenomena including memory consolidation (Girardeau et al., 2009a; Wilson and McNaughton, 1994), mental exploration (Derdikman and Moser, 2010; Gupta et al., 2010), spatial planning (Pfeiffer and Foster, 2013), decision making (Singer et al., 2013), and reinforcement learning (Foster and Knierim, 2012b; Foster and Wilson, 2006). Indeed, each of these proposed roles enjoys some degree of theoretical and empirical support. However, the extent to which these functions correspond to different types of replay is unclear. Our results suggest task demands at the time of replay may dictate the operational mode of replay and thus what is replayed. Namely, during periods of task engagement, when an animal might be expected to be planning or perhaps rehearsing future choices, replay favours adjacent positions and preserves the normal ordering of place cells; consistent with previous work linking replay at decision points with behaviour (e.g. (Diba and Buzsaki, 2007a; Pfeiffer and Foster, 2013; Singer et al., 2013). In contrast, when animals are disengaged from the ongoing task, replay is less focused, reactivating a heterogeneous mixture of positions and paths (e.g. (Davidson et al., 2009; Gupta et al., 2010). Plausibly this latter state, which reflects a composite of recent experiences and engages downstream cortical targets, is likely to be linked with consolidation; a view consistent with previous work (Girardeau et al., 2009b). Furthermore, we showed that successful navigation is predicted specifically by the occurrence of replay focused on the current task. Previous studies have shown performance on spatial memory tasks correlates with the content of replay (Dupret et al., 2010; Papale et al., 2016; Singer et al., 2013), yet we are the first to show that when an animal is actively engaged in a task, *specifically*, the content of replay predicts the accuracy of future spatial decisions. While a causal link is still to be proven, this predictive relationship strengthens the case that different forms of replay perform different functions, indicating that forward local replay is especially important for spatial planning.

In contrast, it is less clear what function the heterogeneous replay occurring during prolonged stationary periods might support. In humans, periods of task disengagement are known to be accompanied by activity in a default mode network (Buckner et al., 2008), which can often incorporate the hippocampus, and is characterised by recall of previous events and autobiographical memories (Greicius et al., 2009; Gupta et al., 2010; Spreng et al., 2009). As such it is tempting to speculate that replay during these periods has a role to play in learning from past experience, potentially being important for memory consolidation (Girardeau et al., 2009b) or could reflect mental exploration of the task environment (Derdikman and Moser, 2010; Gupta et al., 2010)

Finally, we also explored the dynamics of the transition between engaged and disengaged periods, showing that it was rapid but by no means immediate upon corner arrival and departure. How might this switch be mediated? The prefronal-cortex (PFC) might possibly play an important role given its interaction with the hippocampus is known to be modulated by task demands (Place et al., 2016). Another candidate region is the medial entorhinal cortex (MEC). Recent experimental work has shown replay can originate independently from MEC and hippocampus (O’Neill et al., 2017). In turn, replay of MEC grid cells has been suggested to play a role in navigation(Bush et al., 2015; Erdem and Hasselmo, 2012; Erdem and Hasselmo, 2014), while replay originating from the hippocampus seems more likely to contribute to mnemonic processes. As such we propose that during engaged periods, task focused replay is initiated in the MEC but ultimately incorporates place cells. Conversely, during quiescence, replay is mainly hippocampal initiated, and as such reflects the diverse range of trajectories that animals have recently experienced.

## Author Contributions

F.O. and C.B. conceived of the project jointly. F.O. and F.C. performed surgeries. F.O. carried out experiments. F.O. and C.B. performed the analyses. All authors wrote the manuscript.

## Acknowledgements

This work was supported by the Wellcome Trust and the Royal Society. The authors thank Robin Hayman, Francesca Cacucci, and Daniel Bush for comments on the manuscript.

## STAR Methods

### Animals and surgery

Eight male Lister Hooded rats were used in this study. All procedures were approved by the UK Home Office, subject to the restrictions and provisions contained in the Animals Scientific Procedures Act of 1986. All rats (330-400g/13-18weeks old at implantation) received two microdrives, each carrying eight tetrodes of twisted 17μm HM-L coated platinum iridium wire (90% and 10%, respectively; California Fine Wire), targeted to the right CA1 (ML: 2.2mm, AP: 3.8mm posterior to Bregma) and left medial entorhinal cortex (MEC) (ML = 4.5mm, AP = 0.3-0.7 anterior to the transverse sinus, angled between 8-10^<u>o</u>^). Wires were platinum plated to reduce impedance to 200-300kΩ at 1kHz. After rats had recovered from surgery they were maintained at 90% of free-feeding weight with *ad libitum* access to water, and were housed individually on a 12-hr light/dark cycle.

### Recording

Screening was performed post-surgically after a 1-week recovery period. An Axona recording system (Axona Ltd., St Albans, UK) was used to acquire the single-units and positional data (for details of the recording system and basic recording protocol see Barry et al(2007). The position and head direction of the animals was inferred using an overhead video camera to record the location of two light-emitting diodes (LEDs) mounted on the animals’ head-stages (50Hz). Tetrodes were gradually advanced in 62.5um steps across days until place cells (CA1) and grid cells (MEC) were identified.

### Experimental apparatus and protocol

The experiment was run during the animals’ light period to encourage quiet restfulness during the rest session. Animals ran on a Z-shaped track, elevated 75cm off the ground with 10cm wide runways. The two parallel tracks of the Z (190cm each) were connected by a diagonal section (220cm). The entire track was surrounded by plain black curtains with no distal cues. During each track session, animals were required to complete laps on the elevated Z-track. Specifically, the animals were required to run from the start of Arm1 to the end of Arm3, stopping at the track corners and ends in order to receive a food reward. If the animals made a wrong turn at the corners, reward was withheld. Four animals (R2142, R2192, R2198, and R2217) were trained to run on the track for 3 days before recording commenced. For the other animals (R2242, R2335, R2336, R2337), recordings were made from the first day of exposure to the Z-track task.

Following the track session, rats were placed in the rest enclosure for 90 minutes. The rest enclosure consisted of a cylindrically shaped environment (18cm diameter, 61cm high) with a towel placed at the bottom and was located outside of the curtains which surrounded the Z-track. Animals were not able to see the surrounding room while in the rest enclosure. Prior to the experiment, rats had been familiarised with the rest enclosure for at least 7 days. Animals R2242, R2335, R2336 and R2337, were also placed in the rest enclosure for 90 minutes prior to the first Z-track session on day 1 of the experiment. Recordings from this ‘pre-rest’ session were not analysed as part of this study. Following the rest session, animals completed a 20min foraging session in an open field environment. This session was included to enable functional classification of MEC cells and was not analysed in the current study.

### Behavioural Task Performance

To index behavioural fluency we used two measures. First, we recorded the number of incorrect turns animals made at the corners of the track in each trial. Second, we calculated the mean time animals took to complete each lap (total trial duration / number of laps). Learning across days was assessed by correlating these behavioural measures with the number of days of experience the animal had had on the track.

### Data inclusion/exclusion

Sessions where the median decoding error during track running (see ‘Data Analysis’ section below for details) did not exceed 30cm (5% of the length of the track) were included for analyses. Applying this criterion, 10 sessions were excluded. A further two sessions (day4 from R2336, day 8 from R2198) were excluded because of data loss due to the headstages becoming disconnected from the microdrives during the rest session. In total 24 sessions were submitted for further analysis. Only reactivation/replay trajectory events occurring at the two corners were analysed, as the corners represented locations where the animal had a choice as to which way to turn.

### Data Analysis

Ratemaps for the Z-track were generated after first excluding areas in which the animals regularly performed non-perambulatory behaviours (e.g. eating, grooming); the final 10cm at either end of the track and 5cm around each of the two corners. Similarly, periods when the animals’ running speed was <10cm/s were also excluded. To generate ratemaps, the animals’ paths were linearised, dwell time and spikes binned into 2cm bins and smoothed with a Gaussian kernel (σ = 5bins), firing rates were calculated for each bin by dividing spike number by dwell time. Separate ratemaps were generated for runs in the outbound and inbound directions (mean correlation between inbound and outbound ratemaps: r = -0.0032, 1-sample test: p = 0.74). To identify place fields, spatial bins whose rate exceeded the mean firing rate of the cell on the track were only considered. Hippocampal cells were classified as place cells if they exhibited firing greater than its mean rate for 20 contiguous bins and if the peak firing rate was >1hz. Interneurons, identified by narrow waveforms and high firing rates, were excluded from all analyses. Grid cells were identified using a method adopted previously (Olafsdottir et al., 2016b).

Two approaches were employed to analyse place cell reactivations occurring at the two corners (event occurring elsewhere were not analysed). The ‘reactivation’ analysis was used to identify which of the arms making up the track was most strongly represented during events, and the replay ‘trajectory’ analysis identified reactivations of previously taken paths. The methods overlapped significantly for the two approaches, the main difference being the way we carried out the position decoding.

Putative reactivation events were identified based on the activity of hippocampal place cells using a similar method to Pfeiffer and Foster (2013) and Olafsdottir et al(2016b). To identify reactivation events, multi-unit (MU) activity from CA1 place cells were binned into 1ms temporal bins and smoothed with a Guassian kernel (σ = 5ms). Periods when the MU activity exceeded the mean rate by 3 standard deviations were identified as candidate reactivation events. The start and end points of each candidate event were determined as the time when the MU activity fell back to the mean. Events less than 40ms long were rejected. Further, events were excluded if the animals’ movement speed during the event exceeded 3cm/s or if the animals were located away from the two corners (total number of events = 4425). Event detection for the replay trajectory analysis was identical to that for the arm reactivation analysis except we included an additional cell activity criteria for selecting events. Namely, at least 15% of the place cell ensemble or more than 5 place cells, whichever was higher, needed to be active during an event for it be included for analyses.

For position decoding of reactivations, we summed the number of spikes per cell for each event, and a Bayesian framework (Davidson et al., 2009; Olafsdottir et al., 2016b; Zhang et al., 1998) was used to calculate the probability of the animal’s position in each spatial bin given the observed spikes; the posterior probability matrix. Note, two posterior probability matrices were generated for each event, one for inbound runs and one for outbound runs. We then summed the probabilities for each arm (a total of 6 arms given the outbound and inbound runs were decoded separately), to identify which arm was being expressed most strongly during the events. A similar framework was used for the position decoding of replay trajectory events except the spike data was divided up into 10ms temporal bins, and decoding was carried out on each bin separately.

To score the extent to which putative trajectory events represented a constant speed trajectory along the linearised Z-track we applied a line-fitting algorithm (Olafsdottir et al., 2016b). Lines were defined with a gradient (*V*) and intercept (*c*), equivalent to the velocity and starting location of the trajectory. The goodness of fit of a given line (*R*(*V*, *c*)) was defined as the proportion of the probability distribution that lay within 30cm of it. Specifically, where *P* is the probability matrix:

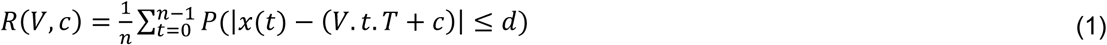

where *t* indexes the time bins of width *T* and *d is* set to 30cm. *R*(*V*, *c*) was maximised using an exhaustive search to test all combinations of *V* between -50m/s and 50m/s in 0.5m/s increments (excluding slow trajectories with speeds > -2m/s and < 2m/s) and c between -15m and 21m in 0.01m increments.

To assess candidate replay events for significance we carried out a spatial shuffle of the place cell ratemaps. Specifically, each ratemap was ‘rotated’ by shifting it relative to the track by a random number of bins drawn from a flat distribution between 1 and the length of the track minus 1 bin. The ratemap for each cell was rotated independently and in each case trailing bins were wrapped around to ensure an equal number of bins were used for each shuffle. This process was repeated 100 times for each event and for each shuffle we recalculated a goodness of fit measure (as described above). This enabled us to estimate the probability of obtaining a given event by chance. Replay trajectory events were defined as those with an individual p-value below 0.025 (a total of 513 trajectory events). Shuffle and data distributions were compared using a 2-sample Kolmogorov-Smirnov test.

A similar approach was used to decode the animals’ locations during track running, except spikes were binned into 500ms temporal bins and location was decoded from the posterior probability matrix using a simple maximum likelihood method. Within each temporal bin an animal’s location was decoded to the bin with the highest posterior probability and this was compared with the known true location (median decoding error for all sessions = 17.5cm, ^~^3% of track length).

### Reactivation Analysis

Corner stops were divided into two temporal sections. The first 5s following an animal’s arrival at the corner and the last 5s preceding an animal’s departure from a corner were categorised as ‘engaged’ periods. Any time interleaving the two was categorised as ‘disengaged’ periods. Reactivation events were analysed separately for the two temporal sections. Temporal sections which lasted <2s were excluded.

For track reactivation events we classified events as ‘congruent’ if the decoded arm belonged to the same run the animal was carrying out at the time of the event. For example, if an animal was carrying out an inbound run and an arm on the inbound run was reactivated. Alternatively, events were classified as ‘incongruent’. Furthermore, reactivation events were considered ‘local’ if they decoded one of the two arms adjacent to the animal. For example, if the animal was located at corner1 during an outbound run, and if arm1 or arm2 were reactivated during this corner period we classified the event as local. To note, for an event to be considered local it also had to be congruent. To estimate the proportion of congruent/local events one would expect to obtain by chance we carried out a cell ID shuffle of each event. Namely, cells active in an events were randomly reassigned to a different cells’s ratemap not active in the event and track decoding carried out on this shuffled data. This process was repeated 100 times and the mean proportion of shuffled congruent/local events used as chance level.

For the replay trajectory analysis, we used the same method to categorise events as congruent or incongruent. However, to categorise local events we estimated the median position during an event and if it lay within 60cm of the animal’s actual position (ahead or behind it) we considered the event to be local. We used an alternative approach to analyse the physical proximity of replay trajectory events. Namely, we calculated the distance between the animal and the start *and* end location of trajectory events. Furthermore, we categorised events as ‘forward’ and ‘reverse’ on the basis of the gradient of the line fit to the place cell posterior probability matrix. Forward events being characterised by a positive gradient which represent reactivation of place cells in the same sequence as they were experienced on the track. Reverse events, characterised by a negative gradient, corresponding to reactivation of place cells in the opposite order to which they would normally be active during track running. To estimate the proportion of congruent/local trajectory events one would expect by chance we randomly reassigned events to either engaged or disengaged periods 100 times, and computed the mean proportion of local and congruent events from the shuffled data. For forward vs reverse classification we used 50% as chance level.

Finally, events preceding correct turns at the corners were categorised as correct events and those preceding incorrect turns error events.

To analyse the temporal dynamics of congruent/local reactivations during corner stops we divided each corner period into 2s time bins, and computed the proportion of congruent/local events for each bin up to 30s following corner arrival/prior to corner departure. For this analysis, we only included corner stops that were at least 30s long. To estimate whether the obtained proportion for each time bin was significantly above chance we binned the data into 5s time bins, advancing in 2.5s increments, and for each bin calculated whether the obtained proportion was significantly greater than chance level.

We carried out a number of control analyses to ascertain the differences obtained for different levels of task engagement could not be accounted for by different LFP states underlying periods associated with task engagement and disengagement. Firstly, to equate the movement speeds during engaged and disengaged events we applied the following approach. Data from the disengaged periods was subsampled to have the same median speed as the engaged periods – this was achieved by removing data corresponding to the highest or lowest speed points first. Second, we excluded replay trajectory events whose best-fit line slope was less than 2m/s (thereby removing replay trajectories that might results from theta phase precession or sequences), following Neill and colleagues (O’Neill et al., 2017). Third, we limited the reactivation events to those whose log(theta/delta) ratio was at least one standard deviation below the mean log(theta/delta) ratio measured during movement (>10cm/s). Fourth, we limited replay trajectory events to those which overlapped with a detected ripple (150-250Hz) event (see details of ripple detection in the subsequent section below).

### Grid Coherence Analysis

To analyse coherence between spatial representations of place and grid cells during hippocampal replay trajectories we followed a method we had adopted previously (Olafsdottir et al., 2016b) – note, this analysis was only conducted for full replay trajectories. Briefly, we applied the same Bayesian framework to the grid cell spikes as we did for the place cell spikes. Hence, for each replay event we also calculated a posterior probability matrix based solely on the observed grid cell spikes. Rather than fitting straight-line trajectories to the periodic grid cell posteriors, we compared the best-fit line from the concurrently recorded place cell posterior. Specifically, we fitted a line with the same intercept and slope as the concurrent place cell event and calculated the proportion of the probability distribution lying within x/2cm of the line. Where x was equal to the average size of the grid cell firing fields recorded from that animal on the linear track. This value we used to index grid-place cell replay coherence. To estimate statistical significance of the observed coherence scores we used the following shuffling procedure: each grid cell posterior was randomly paired with 100 non-concurrent place cell events from the same animal and from the same session. The line fitting procedure to estimate grid-place cell replay coherence, described above, was then re-run on the randomly paired events. If the grid-place cell coherence score obtained in the original data exceeded the 97.5^th^ percentile of its shuffle distribution we deemed the coherence to be better than chance.

To test whether grid-place cell coherence differed between engaged and disengaged periods we carried out the bootstrapping procedure, described in the ‘Statistics’ section below.

### Local Field Potential Analysis

Local field potential (LFP) from CA1 was recorded at 4.8 kHz throughout the experiment. To analyse sharp-wave ripples(Buzsaki et al., 1992) and theta-band (O’Keefe and Nadel, 1978) oscillation the LFP was, firstly, down-sampled to 1.2kHz and then band-pass filtered between 150 and 250Hz and 6-12Hz, respectively. An analytic signal was constructed using the Hilbert transform, taking the form:

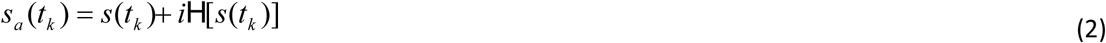

where H specifies the Hilbert transform, *s(t_k_)* is the filtered LFP signal, *t_k_* = *k*Δ, where *k* = indexes the time-step and Δ is the inverse of the sampling rate. An instantaneous measure of power was found by taking the squared complex modulus of the signal at each time point.

For the track-reactivation analysis we down-sampled this measure to 50Hz to match the position sampling rate. To obtain a more stable measure for theta power we computed the ratio between theta and delta (2-4Hz) power for each recording sample and used the log-transformed theta/delta ratio as an estimate of instantaneous theta power(Jackson et al., 2006). For each reactivation event we computed the mean power in the log(ripple/theta) ratio band. For ripple event detection we identified periods where the ripple power exceeded 2.5std of the mean. The start and the end of a ripple event was marked by the point when the power crossed the mean. Events lasting less than 40ms or more than 500ms were excluded and events separated by less than 40ms were joined together. To assess LFP coherence between CA1 and deeper layers of the MEC we used the mscohere function in Matlab 2014a (Mathworks. MA).

R2142 was excluded from the LFP analyses due to problems with EEG recordings.

### Statistics

To assess differences in the proportion of different event types (e.g. congruent events) for engaged and disengaged periods we bootstrapped the data and computed the 95% confidence interval. Namely, we resampled the data with replacement 10,000 times, each time calculating the proportion of a given event type for a particular event period. We then subtracted the proportion of events of a given type occurring during disengaged periods from that occurring during engaged periods, and if 97.5% of the difference scores exceeded 0 we deemed the result significant. To estimate if the obtained proportion significantly differed from chance we counted the number of times the bootstrapped data exceeded the empirically derived chance level (for details of chance calculation see ‘Reactivation analysis’ section above), if more than 97.5% of the bootstrapped data was greater than chance we deemed the data to be significantly above chance. When comparing data and shuffle distributions we used a 2-sample Kolmogorov-Smirnov test. When comparing LFP power and grid-place cell replay coherence during engaged and disengaged periods we carried out the same analysis, but for each bootstrap iteration we computed means rather than proportion. All correlations were carried out using the Pearson product-moment correlation coefficient.

To assess whether there was a significant interaction between task engagement and decision accuracy at the corners we carried out the following analysis. We bootstrapped the data for engaged, disengaged, correct and incorrect events separately, obtaining a bootstrapped distribution of %congruent/%local reactivations for each of the four categories (as described above). For each correct and incorrect bootstrapped distribution pair we computed a difference distribution (by subtracting the correct distribution from the incorrect distribution). We then compared the engaged and disengaged difference distributions to assess whether the engaged difference scores were significantly higher than the disengaged difference scores; implying future decision accuracy modulates the content of place cell reactivations more for reactivations occurring during engaged periods, compared to disengaged periods. If more than 97.5% of the engaged difference scores exceeded the disengaged difference scores we deemed the interaction significant.

### Model based prediction

Binary classification decision trees a simple form of classifier, were used in order to predict, based on the preceding reactivation, if an animal’s next turn would be correct or incorrect. First, because of the disparity in the number of error and correct events, correct events were randomly subsampled to equate the numbers of each class; the procedure was repeated 10 times, model training and testing being completed independently for each iteration. Next, the data used to fit and test each model was segregated further using 10-fold cross validation: reactivation events were randomly divided into 10 equally sized groups, on any given ‘fold’ 9 of these groups were used to train the model and the remaining group used to generate predictions, this process was iterated 10 times such that each events was used once to make and test a prediction. The model was fit using the fitctree function in Matlab 2016b (Mathworks, MA). Three predictor variables based on the content of the replay were used: how congruent and local an event was as well as the average power in the ripple band of the LFP (150-250Hz) during the event. For the model, how congruent and local an event was described by a continuous variable. Specifically, for each event the posterior probability over position for outbound and inbound runs was summed and the summed posterior of the incongruent run subtracted from that of the congruent run; hence, a congruent event would have a positive score. A similar method was used for scoring how local an event was. Namely, the summed posterior of the remote track during an event was subtracted from that of the adjacent tracks. In this instance a local event could either be a congruent or an incongruent event. Predictions were generated by applying the parameters of the trained decision tree to the remaining test data using the Matlab function predict. Prediction accuracy was defined simply as the total proportion of events that were classified correctly.

To establish significance, the same fitting procedure was applied to shuffled data generated by randomly reallocating the response variable relative to the predictor variables; the relationship between the predictors were not changed. This process was repeated 100 times for each of the 10 subsampled iterations, resulting in a distribution of 1,000 prediction accuracies, against which the true prediction accuracy was ranked to generate a p-value.

### Histology

Rats were anaesthetised (4% isoflurane and 4L/min O_2_), injected intra-peritoneal with an overdose of Euthatal (sodium pentobarbital) after which they were transcardially perfused with saline followed by a 4% paraformaldehyde solution (PFA). Brains were carefully removed and stored in PFA which was exchanged for a 4% PFA solution in PBS (phosphate buffered saline) with 20% sucrose 2-3 days prior to sectioning. Subsequently, 40-50μm frozen coronal sections were cut using a cryostat, mounted on gelatine-coated glass slides and stained with cresyl violet. Images of the sections were acquired using an Olympus microscope, Xli digital camera (XL Imaging Ltd.). Sections in which clear tracks from tetrode bundles could be seen were used to confirm CA1 and MEC recording locations.

### Data/code availability

The data that support the findings of this study are available from the corresponding authors upon request.

